# A single-cell multimodal foundation model enables cross modality analysis between transcriptome and chromatin

**DOI:** 10.64898/2026.10.28.762297

**Authors:** Xinyu Bao, Jiajun Zhu, Xiangchen Song, Xiaoshan Zhang, Haoxuan Zeng, Ha T.H. Vu, Haoyun Zhang, Zheyu Zhang, Mingnan Yao, Mengdong Liang, Kai Liu, Stephen CJ Parker, Shuibing Chen, Jie Liu

## Abstract

Most single-cell studies measure either gene expression or chromatin accessibility, and connecting the two usually requires fitting a model to each study. Here we present MUTUAL (MUltiome Transfer in a Unified Aligned Latent space), which keeps the single-cell foundation models scFoundation and EpiAgent frozen, combines their embeddings with linear summaries of each modality and learns a shared RNA– ATAC space from 3.0 million paired cells of 400 public multiome datasets. Applied without refitting to ten datasets not used for training, MUTUAL matched cells’ RNA and ATAC profiles more accurately than methods fitted on other donors of the same dataset, annotated ATAC-only cells more accurately and transferred transcription-factor motif activity to RNA-only cells beyond what cell-type assignment provides. In a maturation series of stem-cell-derived islets, motif activity predicted from RNA largely reproduced the measured stage-wise changes when other libraries of the series served as the reference. MUTUAL makes public multiome data directly usable by single-modality studies.

## 1 Introduction

Gene expression and chromatin accessibility characterize complementary layers of a cell’s regulatory state: accessible chromatin marks the *cis*-regulatory elements and transcription factor (TF) binding sites that are available in a cell, and RNA describes which gene programs are being executed [1].Single-cell atlases now profile each layer at scale, with tens of millions of transcriptomes [2] and more than a million single-cell chromatin accessibility (scATAC-seq) profiles [3, 4], but mostly in separate experiments. Multiome assays, which measure both layers in the same nucleus [5–7], are costlier, and public multiome data amount to only a few million cells, so most cells carry a single modality. ATAC-only cells lack the expression markers on which cell-type annotation usually relies. RNA-only cells lack chromatin information: TF activity can be inferred from the expression of target genes [8, 9], but the accessibility of TF binding motifs measures it more directly [10]. Connecting single-modality data across the two layers is therefore a common task [11]. Many methods integrate unpaired RNA and ATAC data. Some convert accessibility into gene-level activity scores and align the two modalities by canonical correlation analysis or matrix factorization [12, 13]; others learn a joint latent space guided by prior regulatory knowledge or by cells measured in both modalities [14, 15], transfer cell labels across modalities [16], translate one modality into the other [17] or match cells across weakly linked features by optimal transport [18]. These methods differ in what they need from the study at hand. Most are fitted to the cells being integrated; reference-mapping approaches are trained on a reference from the same tissue and then map new samples onto it [19]; and bridge integration maps each single-modality query through one multiome dataset, which must contain the query’s cell types [20]. In each case, what is learned about the correspondence between the modalities stays within one study or one tissue, and accuracy is limited by the data available there.

Pretrained models could carry this knowledge across studies. SCARF [21] pretrains RNA and ATAC encoders and their alignment on 2.7 million cells from 454 public multiome samples and matches cells across modalities without further training, but they learn their encoders on paired data, which remain scarce. Unimodal single-cell foundation models are pretrained on larger or more diverse collections, such as more than 50 million transcriptomes for scFoundation [22] and about 5 million chromatin accessibility profiles for EpiAgent [23]. Each, however, defines its own embedding space, so the RNA and ATAC embeddings of the same cell cannot be compared. In vision–language modeling, contrastive training on paired examples creates shared spaces that transfer to new data without retraining [24], and the encoders of the two modalities can be pretrained separately and kept frozen, with only a small connecting module trained [25]. Whether single-cell foundation models can serve as such frozen encoders for RNA and chromatin, and what they add to simple linear summaries of each modality, has not been systematically tested.

Here we present MUTUAL (MUltiome Transfer in a Unified Aligned Latent space), a model that places RNA-only and ATAC-only cells in one shared embedding without refitting (Fig.1). MUTUAL keeps scFoundation and EpiAgent frozen and describes each cell in three ways: by its foundation-model embedding; by a linear summary computed once on the training data (principal components of expression [12] or latent semantic indexing of accessibility [26]); and by its individual highly variable genes or accessible regulatory elements. Small neural networks for each modality are trained on 3.01 million paired human cells from 400 public multiome samples in 14 tissue groups to bring the two profiles of each cell, and cells of the same annotated type, close together while pushing other cells apart. After training, each modality is embedded independently by a fixed model, so new RNA-only samples, ATAC-only samples and single-modality reference atlases share one space, and labels and chromatin features can be transferred between them. In effect, MUTUAL replaces a tissue-matched bridge dataset with a mapping learned once from hundreds of multiome datasets.

**Fig. 1:**
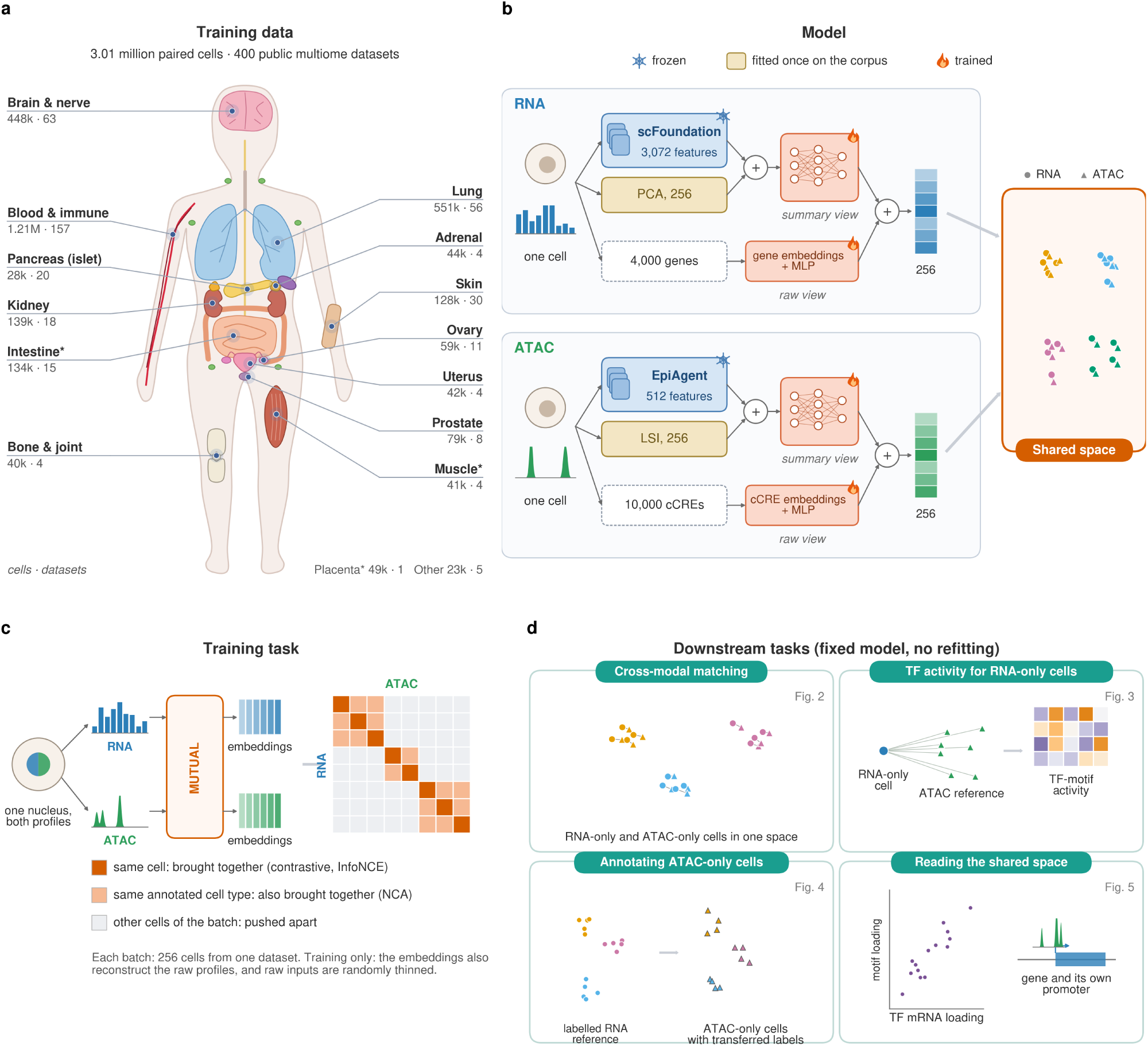
MUTUAL learns a shared RNA–chromatin embedding once from public multiome data. **a**, Training data: 3.01 million paired cells (RNA and ATAC measured in the same nucleus) from 400 public multiome datasets, by tissue of origin, shown on a schematic human body (number of cells *·* number of datasets). Asterisks mark groups made up mostly or entirely of *in vitro* cells (an intestinal culture, cultured myotubes and a trophoblast cell line). **b**, Model. For each modality, a summary view joins the embedding of a frozen foundation model (scFoundation for RNA, EpiAgent for ATAC) with a linear summary fitted once on the training corpus (principal component analysis, PCA, of gene expression; latent semantic indexing, LSI, of chromatin accessibility), and a raw view reads panel genes or candidate *cis*-regulatory elements (cCREs) through trained feature embeddings. Trained networks (multilayer perceptrons, MLPs) map each view to 128 values, and the two are combined into a 256-value embedding in a space shared by both modalities. **c**, Training task. For a batch of 256 cells from one dataset, the RNA and ATAC embeddings of the same cell are drawn together and those of other cells pushed apart (InfoNCE), and cells with the same annotated type are also drawn together (neighborhood components analysis, NCA); reconstruction of the raw profiles and random thinning of raw inputs are used only during training. **d**, Downstream tasks performed with the fixed model: matching RNA and ATAC profiles across modalities (Fig.2), inferring chromatin transcription factor (TF)-motif activity for RNA-only cells from an unpaired ATAC reference (Fig.3), annotating ATAC-only cells from a labeled RNA reference (Fig.4) and examining what the shared space aligns (Fig.5). Body outline and organ positions are schematic.

We evaluated MUTUAL on 10 human multiome datasets that were not used for training, hiding each cell’s RNA–ATAC pairing from every method. Without refitting, MUTUAL matched the RNA and ATAC profiles of the same cells more accurately than SCARF and than GLUE, MultiVI and Seurat bridge integration fitted on other donors of the same dataset (Fig.2). It transferred chromatin TF-motif activity from ATAC references to RNA-only cells beyond what RNA-based inference or cell-type assignment provides, with the highest agreement of the methods tested (Fig.3). In a maturation series of stem-cell-derived islets, its RNA embedding ordered the stages along a trajectory, and motif activity predicted from RNA, with the series’ other libraries as the chromatin reference, largely reproduced the stage-wise changes measured in the series and reported by the original study. It annotated ATAC-only cells from other donors’ labeled RNA more accurately, on average, than these methods, and its vote confidence allowed the most reliable labels to be kept (Fig.4). Finally, its shared space was organized by cell identity but also paired cells within cell types, aligned TF expression with the activity of the TFs’ motifs and linked genes to their own promoters, and the foundation-model inputs added modestly to the linear summaries (Fig.5). MUTUAL thus makes public multiome data directly usable by single-modality studies.

**Fig. 2:**
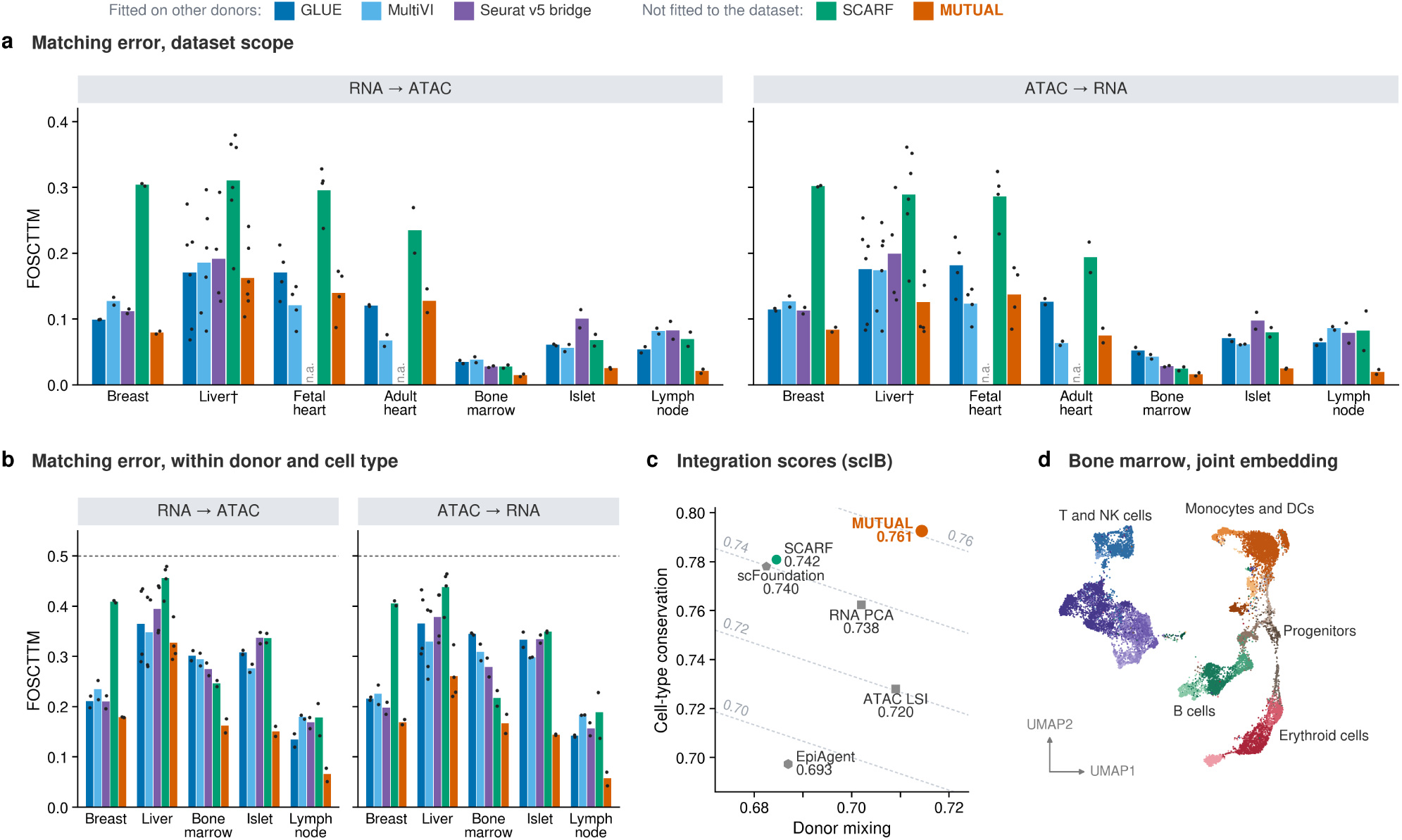
MUTUAL pairs the RNA and ATAC profiles of held-out cells without being fitted to them. **a**, Matching error (FOSCTTM, the fraction of candidate cells ranked above a query’s true partner; 0 is perfect and 0.5 is chance) at the dataset scope, where the candidates are all cells of the scored fold. RNA *→* ATAC, RNA queries ranked against ATAC candidates; ATAC *→* RNA, the reverse. The individuals of each dataset were split into two folds. GLUE and MultiVI were fitted on one fold, and the Seurat v5 bridge mapped cells through that fold’s paired cells (together, fitted on other donors); all methods were then scored on the other fold, and the folds were swapped. MUTUAL and SCARF were not fitted to the evaluation data. Bars, mean over a tissue’s datasets (liver, three datasets; fetal heart, two; other tissues, one) of the mean of the two folds; points, the single folds of each dataset. Ten evaluation datasets (342,608 cells from 76 individuals). The bridge was run only on the datasets with cell-type labels: n.a., not run; *†*, the bridge’s liver bar averages the two labeled liver datasets. **b**, Within-type scope: the candidates are the cells of the query’s own donor and cell type (pools of 64–2,000 cells), on the labeled datasets of **a** (liver, two datasets); dashed line, chance. **c**, scIB integration scores averaged over the six labeled datasets (at most 4,000 cells per dataset, the same cells for every method). MUTUAL and SCARF were scored on each cell’s concatenated RNA and ATAC embeddings, and the single-modality inputs on their own (grey; pentagon and hexagon, the foundation models; squares, RNA principal components (PCA) and ATAC latent semantic indexing (LSI)). Numbers, overall score (0.6*×* conservation + 0.4*×* mixing; Methods); dashed lines, equal overall score. **d**, Uniform manifold approximation and projection (UMAP) of the joint MUTUAL embedding of 18,305 bone-marrow cells from 13 samples of 10 individuals, colored by the dataset providers’ 22 cell types in one color family per lineage (NK, natural killer; DC, dendritic cell). In **a**, **b** and **d**, MUTUAL is the ensemble of its three models; in **c**, its scores are means over the three models.

**Fig. 3:**
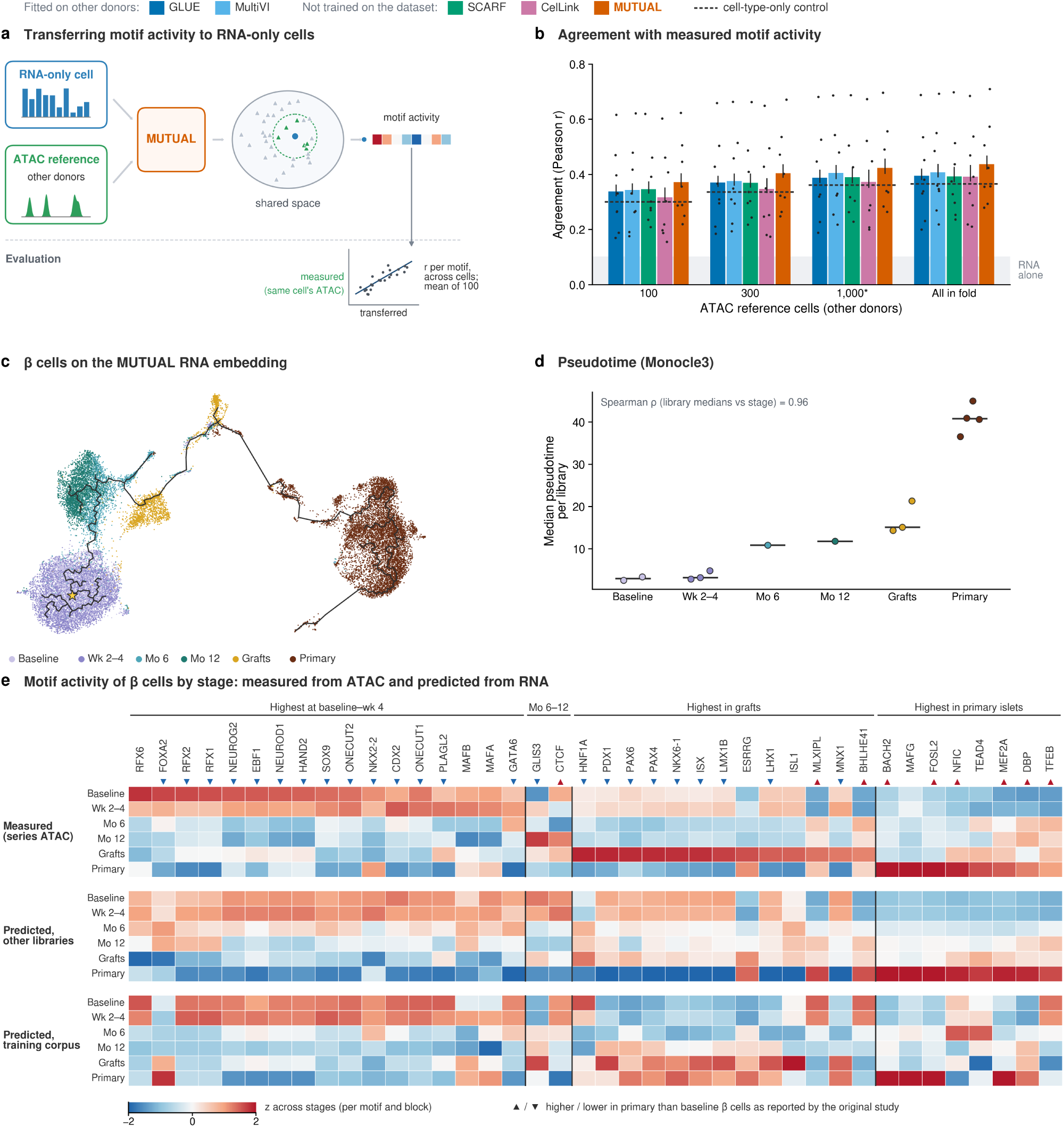
MUTUAL transfers TF-motif activity to RNA-only cells from an ATAC reference. **a**, Each RNA-only cell receives the similarity-weighted mean chromVAR motif activity of its 20 nearest ATAC reference cells (other donors). Agreement with the activity measured from the cell’s own ATAC, hidden during transfer, is the Pearson correlation across cells, per motif, averaged over motifs. **b**, Agreement over all scored cells of the eight datasets from tissues absent from training (100 most variable motifs), against the number of reference cells. GLUE and MultiVI were fitted on the other fold of donors; SCARF and CelLink were not trained on these data. **c**–**e**, Stem-cell-derived islets (14 libraries; excluded from training). **c**, UMAP of the MUTUAL RNA embedding of 23,576 *β* cells coloured by stage (Wk, weeks; Mo, months in culture; grafts, recovered after transplantation into mice), with the Monocle3 principal graph (star: root). **d**, Median pseudotime per library (dots) and stage medians (lines). **e**, *β*-cell motif activity by stage for 40 motifs listed before any prediction, measured from the series’ ATAC (top) or predicted from RNA with the other libraries (middle) or the 3.0-million-cell training corpus (bottom) as the ATAC bank; stage means z-scored per motif and block. Triangles, higher (▴)_8_ or lower (▾) in primary than in baseline *β* cells as reported in 30.

**Fig. 4:**
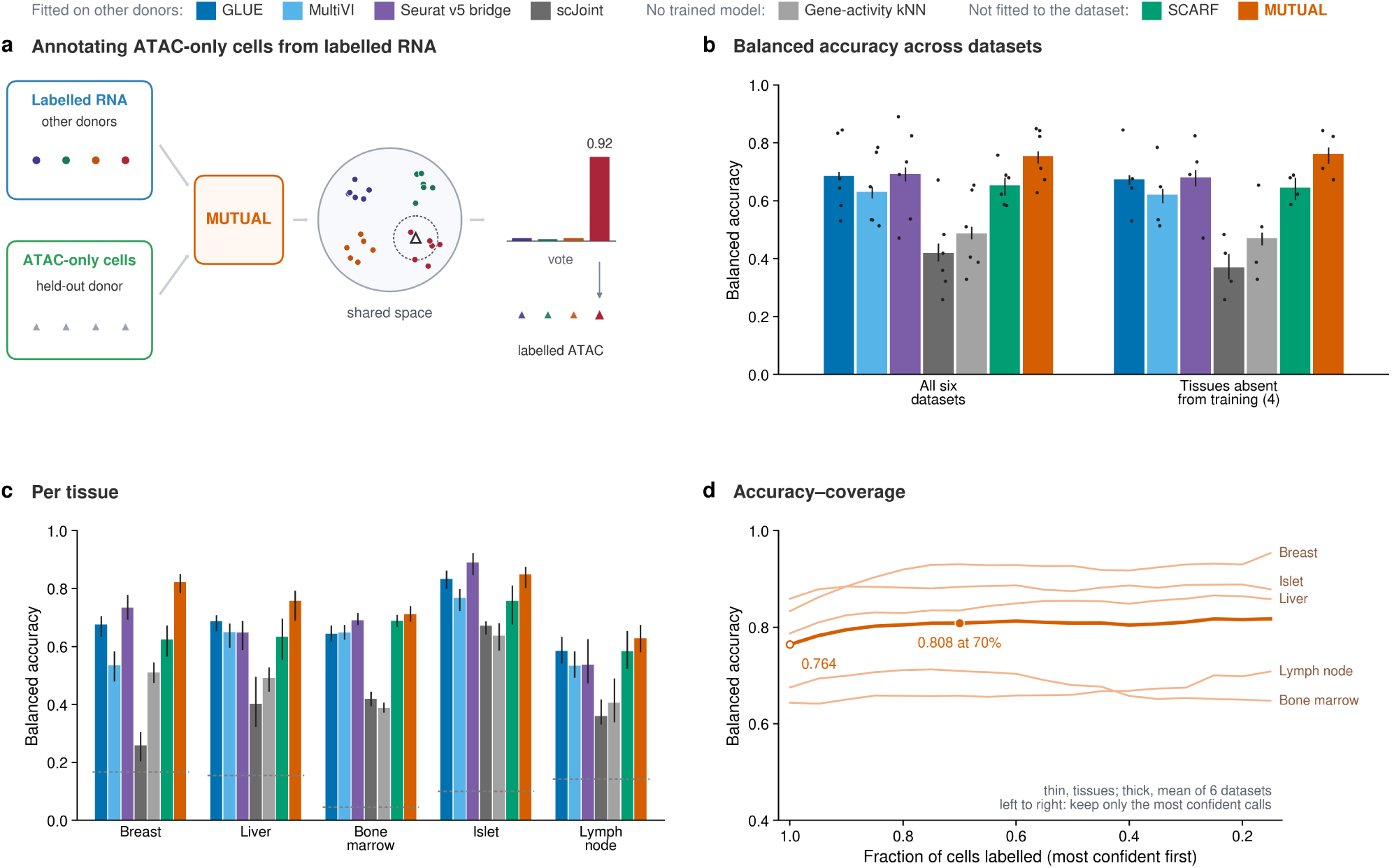
MUTUAL annotates ATAC-only cells from labelled RNA of other donors without training on the target data. **a**, Workflow. The ATAC profiles of a held-out donor and the labeled RNA of other donors are embedded by the fixed model and centered per donor; each ATAC cell takes the similarity-weighted vote of its 15 nearest RNA cells (nearest-neighbor vote), and the winning share of the vote (here 0.92) is its confidence. Transferred labels are scored against the reference labels of the held-out donor’s cells. **b**, Balanced accuracy (mean recall over cell types) in the fold design, averaged over the six labeled datasets and over the four labeled datasets from tissues absent from training. Bars, means; whiskers, 95% CI from donor-block bootstrap; points, single datasets. GLUE, MultiVI and scJoint were fitted on the other fold of donors, and the Seurat v5 bridge mapped queries through that fold’s paired cells; MUTUAL and SCARF were not fitted to the target data, and the gene-activity nearest-neighbor (kNN) baseline uses no trained model. MUTUAL is the mean of its three models. **c**, Balanced accuracy per tissue (liver, mean of the two labeled liver datasets); whiskers, 95% CI; dashed segments, chance (one over the number of cell types). Labels are from the original authors or providers. **d**, Accuracy–coverage of MUTUAL in the all-donor reference design: balanced accuracy among the cells labeled with the highest confidence; from left to right, only the cells with the most confident votes are kept (x axis, fraction of cells labeled). Thin lines, tissues (liver, mean of the two liver datasets); thick line, mean of the six datasets.

**Fig. 5:**
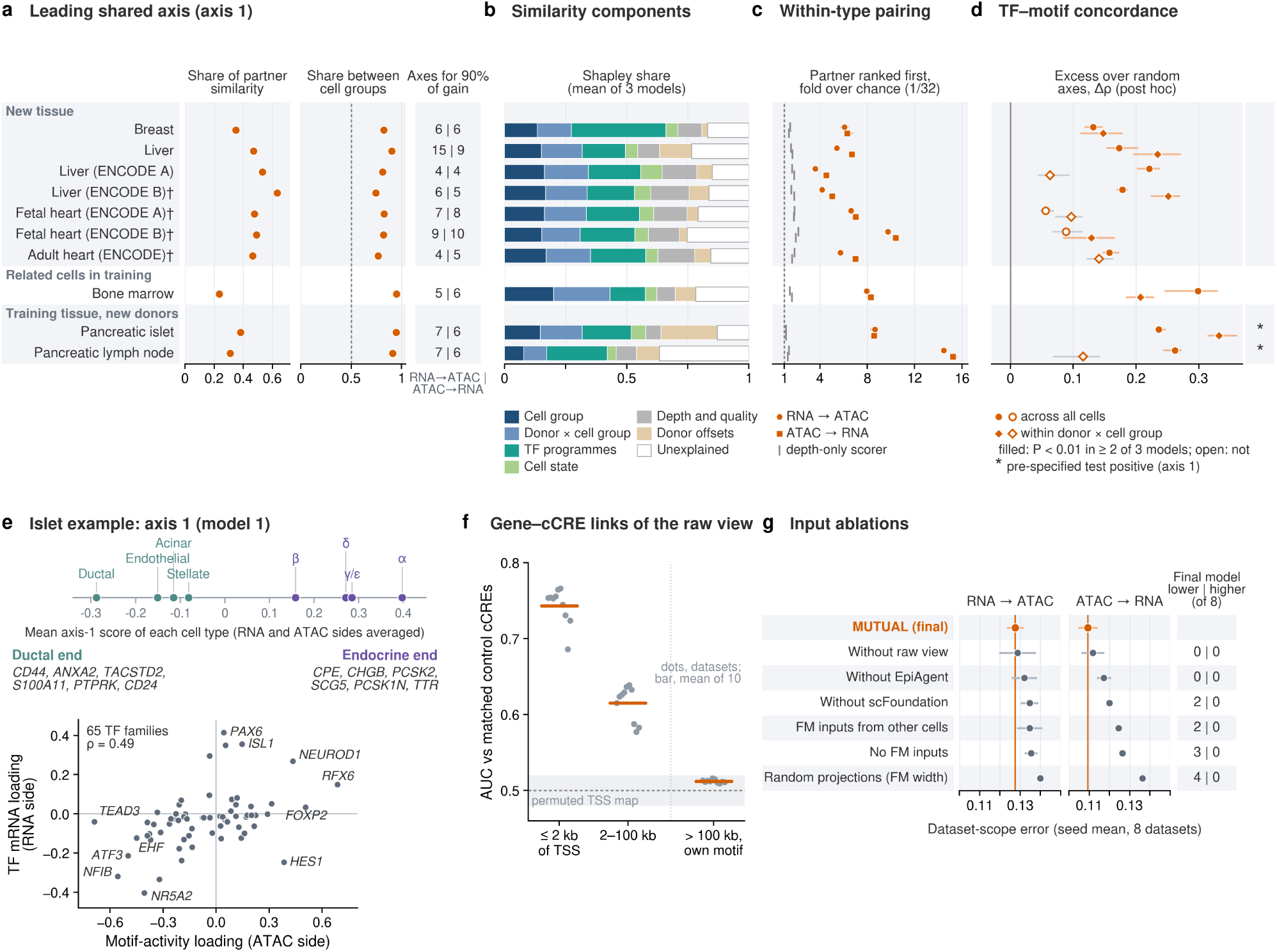
What the shared space aligns and what its inputs contribute. **a**, Shared axes are the eigenvectors of the symmetrized cross-covariance between RNA and ATAC embeddings, estimated on one half of each dataset’s cells and evaluated on the other. Left, share of the mean similarity between partners carried by the leading axis (axis 1); middle, share of the variation along axis 1 that lies between cell groups (axis 1 is called an identity axis right of the dashed line); right, number of leading axes needed to recover 90% of the matching gain over chance in each query direction (median of three models). **b**, Shapley decomposition of the similarity between partners (at most 5,000 cells per dataset; mean of three models). **c**, Fold over chance (1/32) with which the true partner is ranked first in pools of 32 cells of one cell group and donor (569 strata in total, 200 pools each); grey ticks, a depth-only scorer, one per query direction; dashed line, chance, which the permutation floor matched. **d**, Agreement between the loadings of TF mRNA and those of the same TF families’ motif activity on the five leading axes, shown as the excess of its Spearman correlation over that of random shared axes (a test defined after the first results had been seen), with loadings computed across all cells (circles) or within donor *×* cell group (diamonds). Filled, *P <* 0.01 in at least two of three models; open, not; because each dataset has its own null spread, significance need not follow the size of the excess. Asterisks, datasets in which the pre-specified test was informative and positive for axis 1. **e**, Islet example. Top, mean axis-1 score of each cell type with at least 20 cells (average of the RNA-side and ATAC-side scores); middle, the six genes at each end whose expression correlates most strongly with the RNA-side score within donors; bottom, loadings of motif activity and of TF mRNA on axis 1 for all 65 TF families (family means; labels, each family’s TF with the strongest mRNA loading). **f**, Coupling of raw-view genes to raw-view cCREs, measured from gradients of the cross-modal similarity, as the area under the curve (AUC) for cCREs within 2 kb of the gene’s transcription start site (TSS), 2–100 kb away, or more than 100 kb away and carrying the motif of the TF that the gene encodes, each against 20 matched control cCREs of the same gene (2,486–2,605 genes per dataset within 2 kb; mean of three models); shaded band, 0.5 *±* 0.02, the pre-specified tolerance for the permuted TSS map (observed AUCs with the map permuted, 0.489–0.514). **g**, Input ablations: mean dataset-scope matching error over the eight datasets from tissues absent from training, scored on all their cells (mean *±* s.d. of three models; orange line, final model). Right, number of these datasets on which the final model had a lower or a higher

## 2 Results

### 2.1 A shared RNA–ATAC embedding learned once from public multiome data

MUTUAL learns the correspondence between gene expression and chromatin accessibility once, from public paired data, and then applies it to new single-modality data without refitting. Each modality is described in two complementary views (Fig.1b). The summary view joins a frozen foundation-model embedding, from scFoundation for RNA and EpiAgent for ATAC, with a linear summary of the same profile that is computed once on the training corpus (principal components of expression or latent semantic indexing of accessibility). The raw view reads 4,000 variable genes or 10,000 frequently accessible candidate *cis*-regulatory elements (cCREs) directly through learned feature embeddings. Small networks map both views to a single 256-dimensional embedding per modality, so that RNA and ATAC profiles become directly comparable.

The networks are trained on cells whose RNA and ATAC profiles were measured in the same nucleus (Fig.1c). Within each batch of 256 cells, a contrastive objective [24] draws the two profiles of the same cell together and pushes those of other cells apart, and a neighbourhood-components term [27] additionally draws together cells with the same annotated type; most of these annotations were assigned automatically (Methods). During training only, the embeddings are also asked to reconstruct the raw profiles, and the raw inputs are randomly thinned. Because the foundation models stay frozen, 9.8 million parameters are trained (6.2 million are used at inference), and a model trains in 49 min on one GPU once the foundation-model features have been computed; we trained three models with different random seeds.

The training corpus comprises 3.01 million paired cells from 400 public multiome datasets spanning 14 tissue groups (Fig.1a). Blood and immune cells account for 40% of the cells, followed by lung (18%) and brain and nerve (15%), and some groups consist mostly of cells grown *in vitro*. Because each batch is drawn from a single dataset and datasets are sampled uniformly, small datasets carry more weight in training than their share of cells.

Once trained, the model embeds each modality on its own: an RNA-only sample, an ATAC-only sample or a single-modality reference atlas enters the shared space without refitting (Fig.1d). We used this property for four tasks: matching RNA and ATAC profiles across modalities (Fig.2), inferring chromatin transcription factor (TF)-motif activity for RNA-only cells from an unpaired ATAC reference (Fig.3), annotating ATAC-only cells from labelled RNA references (Fig.4) and examining what the shared space aligns (Fig.5). We evaluated these tasks on 10 multiome datasets with 342,608 cells from 76 individuals that were held out of training: eight from tissues without training cells (breast, liver and heart, bone marrow) and two from new donors of training tissues (pancreatic islet and lymph node). To compare fairly, we fitted GLUE and MultiVI on the other donors of each evaluation dataset and scored every method on the held-out donors. The pairing of the scored cells’ RNA and ATAC profiles was hidden from every method and used only for scoring.

### 2.2 MUTUAL pairs the RNA and ATAC profiles of held-out cells without being fitted to them

We first asked how well the shared embedding pairs the two profiles of cells that were held out of training. For each RNA profile we ranked the ATAC profiles of all candidate cells, including its own, by their similarity to it, and recorded the matching error: the fraction of other cells ranked above the true partner (FOSCTTM, the fraction of samples closer than the true match [28]; 0 is perfect and 0.5 is chance). ATAC profiles were scored against RNA in the same way, and the two query directions are reported separately (as RNA to ATAC / ATAC to RNA). To compare MUTUAL with methods that must be fitted to each dataset, we split the individuals of every evaluation dataset into a fitting fold and a scored fold, scored all methods on the scored fold and then swapped the two. GLUE [14] and MultiVI [15] were trained on the fitting fold, and the Seurat v5 bridge integration [20], adapted for matching (Methods), mapped both modalities of the scored cells through the fitting fold’s paired cells; we refer to these three as fitted on other donors. MUTUAL and SCARF, [21] a pretrained RNA–ATAC model applied with its released weights, were not fitted to the evaluation data. The candidates were all cells of the scored fold (dataset scope, primary), the cells of the query’s donor or the cells of the query’s donor and cell type (within-type scope). Unless stated otherwise, MUTUAL denotes the ensemble of three models trained with different random seeds, whose embeddings are concatenated (Methods).

Fig.2a shows the ten evaluation datasets, with the three liver datasets and the two fetal heart datasets averaged per tissue. Across them, MUTUAL’s mean dataset-scope error was 0.104 / 0.087, against 0.122 / 0.132 for GLUE, 0.117 / 0.115 for MultiVI and 0.223 / 0.212 for SCARF. MUTUAL was more accurate than GLUE on eight of the ten datasets and than MultiVI on six, and less accurate on one (MultiVI on the adult heart). It was more accurate than the bridge on five of the six labeled datasets on which the bridge was run and not different on the sixth, a liver dataset (mean 0.067 / 0.058, against 0.118 / 0.120), and its error was lower than SCARF’s in both directions on all ten datasets. The largest differences were in tissues represented in training. On the new donors of the pancreatic islet and pancreatic lymph node, whose other donors are in the training data, MUTUAL’s error was 23–45% of that of the methods fitted on other donors, and on bone marrow, whose cell types are related to the blood and progenitor cells in training, it was 31–57%. Among the seven datasets from tissues absent from training, MUTUAL was more accurate than GLUE on five, with the other two comparable, and MUTUAL was more accurate than MultiVI on three: breast and two of the three liver datasets.

At the within-type scope (Fig.2b), the candidates share the query’s donor and cell type, so cell type cannot help and matching must rely on differences between individual cells. Here MUTUAL had a lower error than every method fitted on other donors in both query directions on five of the six labeled datasets, for example 0.151 / 0.144 against 0.276–0.337 on the islet and 0.066 / 0.058 against 0.135–0.183 on the pancreatic lymph node. On one of the two labeled liver datasets, GLUE and MultiVI had lower errors from RNA to ATAC and higher errors from ATAC to RNA. Every method matched better than chance on every dataset, although SCARF approached chance on one liver dataset (0.476 / 0.462).

Finally, we scored the joint representation of paired cells, in which each cell’s RNA and ATAC embeddings are concatenated, with the scIB metrics [29], which reward keeping cell types apart (conservation) while mixing donors (mixing). This measures how well a representation preserves cell-type structure across donors; it does not test where single-modality cells are placed. Averaged over the six labeled datasets, MUTUAL scored 0.761 overall (mean of its three models), above SCARF (0.742, also scored on concatenated embeddings). It also scored above each single-modality input scored on its own (scFoundation, EpiAgent, RNA PCA and ATAC LSI; 0.693–0.740), which is a reference point rather than a like-for-like comparison, as each input carries one modality. It had the highest conservation (0.793) and mixing (0.714) scores of these six representations (Fig.2c). Dataset by dataset, MUTUAL scored higher than EpiAgent and ATAC LSI on all six datasets and higher than SCARF, scFoundation and RNA PCA on five; it scored lower only on breast, against SCARF and RNA PCA. In bone marrow, the joint embedding resolved the major lineages, with progenitor cells, including haematopoietic stem cells, at the junction of the myeloid, erythroid and B-cell branches (Fig.2d).

### 2.3 MUTUAL transfers TF-motif activity to RNA-only cells from an ATAC reference

An RNA-only cell placed in the shared space has ATAC neighbors, so features measured in those neighbors can be transferred to it (Fig.3a). We transferred motif activity, the chromVAR deviation score of each TF motif [10]: each RNA-only query received the similarity-weighted mean motif activity of its 20 nearest cells in an ATAC reference from other donors, whose RNA profiles were not used. Because the evaluation datasets are multiome, the query’s own ATAC profile, hidden during transfer, gives the measured activity. For each motif we computed the Pearson correlation between transferred and measured activity across all scored cells of a dataset, after adjusting for quality covariates, and averaged it over the 100 motifs that vary most across those cells. As in Fig.2, the donors of each dataset were split into two folds; queries and references came from different donors of one fold, GLUE and MultiVI were fitted on the other fold, SCARF was used with its released weights, and CelLink [18], which solves an optimal-transport plan between the query and reference cells, was not trained. All methods used the same queries, references and transfer procedure, on embeddings centered per donor (Methods). Shuffled controls gave *|r| <* 0.001, and TF activity estimated from the queries’ own RNA reached at most 0.10 on average.

Across the eight datasets from tissues absent from training, MUTUAL had the highest mean agreement at every reference size, from 0.372 with 100 reference cells to 0.437 with all reference cells of the fold, against 0.317–0.408 for the other methods (Fig.3b). MUTUAL’s lead over the strongest of MultiVI, GLUE and SCARF was 0.019–0.029 across reference sizes. Its lead over CelLink was 0.045–0.057.

We next applied MUTUAL to a maturation series of stem-cell-derived islets (SC-islets) [30] that was excluded from training. The series comprises 14 multiome libraries (97,247 cells): newly differentiated (baseline) SC-islets, SC-islets after 2–4 weeks and 6 and 12 months in culture, grafts recovered 6 months after transplantation into mice, and primary human islets; each time point in culture is a single library. We embedded the RNA profiles of 23,576 *β* cells and ordered them with Monocle3 [31], with a single principal graph over all cells (Methods). On the embedding, *β* cells of the stem-cell-derived libraries and of primary islets formed separate groups, which the graph connects through a sparse region (Fig.3c). With the root at the tip of the graph richest in baseline cells, pseudotime increased with stage: library medians were 2.5 and 3.4 at baseline, 2.9–4.8 after 2–4 weeks, 10.9 and 11.8 at 6 and 12 months, 14.3–21.3 in grafts and 36.5–45.0 in primary islets (Spearman *ρ* = 0.96 between library medians and stage order over the 14 libraries; Fig.3d). This ordering was similar to that of the same pipeline run on library-corrected principal components (*ρ* = 0.96). Within libraries it did not follow the score (*ρ* = 0.03), so the ordering reflects differences between libraries, which coincide with stages.

We then predicted motif activity for every RNA profile of the series from two ATAC references, or banks: the other libraries of the series (excluding each cell’s own library, so that the single libraries of months 6 and 12 had no same-stage cells in the bank) and the 3.0 million ATAC profiles of our training corpus, which contains no stem-cell-derived cells. Before computing any prediction, we listed 40 TF motifs, 36 of which the original study reports to change between these stages (Methods). In the series’ own ATAC, these motifs separated the stages in *β* cells (Fig.3e): motifs of RFX, NEUROG2, NEUROD1, HAND2, SOX9 and ONECUT factors were most active at baseline and after 2–4 weeks, homeobox motifs such as those of PDX1, NKX6-1 and PAX4 in grafts, and motifs of AP-1 (FOSL2), BACH2, MAFG, NFIC, MEF2A and DBP in primary islets; these measured directions matched 84% of the directions reported in the original study. With the other libraries as the bank, the predicted stage profiles agreed with the measured ones (per-motif correlation of predicted with measured library means, median 0.74 over the 40 motifs; the primary-versus-baseline direction matched for 39 of 40 motifs), although the rise of homeobox motifs in grafts was missed. With the training corpus as the bank, agreement was lower (median 0.53; direction matched for 24 of 40 motifs, against 20 expected by chance). AP-1, BACH2 and DBP motifs were again highest in primary islets, ONECUT, NEUROD1, NEUROG2 and HAND2 motifs at baseline, and homeobox motifs rose in grafts, consistent with the original study, which reports higher AP-1 and BACH2 and lower ONECUT2, RFX1 and NEUROD1 motif activity in primary than in stem-cell-derived *β* cells. However, homeobox and FOXA2 motifs were also predicted high in primary islets, opposite to the measurements, and the 30 motifs that best separate the libraries in the measured data were not recovered from the corpus (median correlation 0.07): the nearest corpus neighbors of early stem-cell-derived *β* cells were mostly enteroendocrine cells, because the corpus contains no chromatin of this cell state. Thus MUTUAL gives RNA-only samples the stage-specific chromatin motif activity of a related reference and, from public multiome data alone, part of the reported changes, chiefly those that separate primary from stem-cell-derived *β* cells.

### 2.4 MUTUAL annotates ATAC-only cells from labeled RNA of other donors without training on the target data

Because RNA and ATAC profiles share one space, the cell-type labels of an annotated RNA reference can be carried to unlabeled ATAC-only cells by a nearest-neighbor vote (Fig.4a). In each labeled evaluation dataset we held out one donor at a time, embedded its ATAC profiles as queries and labeled each query

by a similarity-weighted vote of its 15 nearest RNA cells from other donors. We scored balanced accuracy, the mean recall over cell types, against each dataset’s reference labels (Methods), counting queries whose type had at least five reference cells. In the fold design, as in Fig.2, the references were the other donors of the query’s fold, and methods that are fitted to target data used only the other fold: GLUE, MultiVI, scJoint [16] and the Seurat v5 bridge integration [20], which maps queries through up to 10,000 paired cells of the other fold. scJoint was trained on the reference RNA and the other fold’s ATAC profiles. SCARF and a gene-activity nearest-neighbour (kNN) baseline, which uses no trained model, were also scored.

Averaged over the six labeled datasets, MUTUAL reached a balanced accuracy of 0.755 (95% CI 0.729– 0.771; 0.767 for the three-model ensemble), higher than the Seurat v5 bridge (0.692; paired difference +0.064 on the same bootstrap draws, CI +0.048 to +0.073) and GLUE (0.686; +0.069, CI +0.048 to +0.082), and higher than MultiVI (0.631), SCARF (0.653), scJoint (0.419) and the gene-activity kNN baseline (0.487; Fig.4b). On the four labeled datasets from tissues absent from training, MUTUAL reached 0.763, against 0.680 for the bridge, 0.674 for GLUE and 0.621 for MultiVI. Per dataset (Fig.4c shows the two liver datasets averaged), MUTUAL was more accurate than GLUE on breast, bone marrow and one liver dataset and showed no difference from it on the other three; it was more accurate than the bridge on breast, the pancreatic lymph node and one liver dataset, showed no difference on the other liver dataset and bone marrow, and was less accurate on the pancreatic islet. Also, it was more accurate than MultiVI, SCARF and the gene-activity kNN baseline on every dataset. GLUE and MultiVI fitted on all cells of each dataset, MultiVI also using the scored pairs, reached 0.737 and 0.723.

The vote also gives each label a confidence: labeling only the 70% of cells with the most confident votes raised MUTUAL’s balanced accuracy from 0.764 to 0.808 (Fig.4d).

### 2.5 The shared space is organized by cell identity and also carries TF-program and gene–promoter signals

To identify what the matching relies on, we decomposed the similarity between each cell’s RNA and ATAC embeddings into shared axes: the eigenvectors of their symmetrized cross-covariance, estimated on one half of each dataset’s cells and evaluated on the other half, separately for each of the three models (Methods). A few axes carried most of the match. The leading axis, axis 1, carried 24–63% of the mean similarity between partners, and 4–15 axes recovered 90% of the matching gain over chance, that is, of the reduction in matching error from 0.5 to that of the full embedding (Fig.5a). In every dataset and model, at least 73% of the variation along axis 1 lay between cell groups (author-annotated cell types, or clusters of RNA profiles in the four datasets without labels), so axis 1 was an identity axis in every dataset.

We then attributed the similarity between partners to properties of the cells with Shapley values [32]. The cell group was credited with 8–20% of it and donor-specific differences within cell groups (donor *×* cell group) with a further 9–23%, TF programs (leading principal components of TF-motif activity and of TF mRNA) with 14–39%, read depth and quality metrics with 6–16% and other cell-state programs with 4–9% (Fig.5b). Donor offsets, the differences between donors’ mean embeddings that both modalities share, accounted for 2–23%, most in the islet, and 13–37% was not attributed to these properties.

Cell identity did not account for the whole match. In pools of 32 cells of the same cell group and donor, the true partner was ranked first for 11–48% of queries, 3.6–15.2 times as often as expected by chance (Fig.5c). This exceeded a permutation floor, which matched chance, and a depth-only scorer, which pairs cells by their numbers of detected genes and accessible cCREs alone (1.1–2.2 times chance), in all 10 datasets and in each model. Fixed pools of 32 give every dataset the same chance level, which the within-type matching error of Fig.2b, computed over pools of different sizes, does not.

The model was given no TF–motif pairs. If its shared axes nevertheless follow regulatory programs, the mRNA of an activating TF and the activity of its own motif should load on them with the same sign. For each of the five leading axes, we correlated across TF families the loadings of TF mRNA with those of the families’ motif activity (chromVAR deviations; 49–65 families per dataset). The pre-specified test is interpretable only where its positive control, contrasts between cell groups scored with the same statistic, is detected. This was the case only for the islet and the pancreatic lymph node, and in both, mRNA and motif loadings agreed on axis 1 in each of the three models (Spearman *ρ* = 0.49 over 65 families in the islet and 0.45 over 49 in the lymph node, means of the three models; permutation *P ≤* 0.002). The test’s negative control, the same statistic after permuting cells within donors, gave false positives in five other datasets, so we compared the observed agreement with that of random shared axes instead, a test defined after these results had been seen (Methods). Pooled over the five leading axes, the agreement exceeded that of random axes in 8 of the 10 datasets (*P <* 0.01 in at least two of three models), but not in the two fetal heart datasets (Fig.5d). With the loadings computed within donor *×* cell group, which removes differences between cell groups, the agreement exceeded random axes in 6 datasets, 5 of them among these 8, so there it was not explained by cell identity alone. In the islet, axis 1 ran from ductal, endothelial, acinar and stellate cells to *β*, *δ*, *γ*/*ɛ* and *α* cells, with endocrine genes such as *CHGB*, *PCSK2* and *SCG5* at one end and ductal genes such as *CD44*, *ANXA2* and *TACSTD2* at the other (Fig.5e). Across all 65 TF families, motif-activity and mRNA loadings on this axis were correlated (*ρ* = 0.49 in model 1): the families of RFX6, NEUROD1 and ISL1, which include two TFs required for islet endocrine development [33, 34], loaded on the endocrine end in both modalities, and those of NFIB, NR5A2 and ATF3 on the ductal end.

The raw view reads cCREs through feature embeddings that were initialized at random, and none of the inputs carries genomic coordinates, so the model was never told where a cCRE lies relative to a gene. Using gradients of the cross-modal similarity, we measured how strongly each raw-view gene is coupled to each raw-view cCRE. In all 10 datasets, cCREs within 2 kb of a gene’s own transcription start site (TSS) were coupled to it more strongly than matched control cCREs of the same gene (area under the curve, AUC, 0.69–0.77, mean of the three models; lower 95% confidence bound above 0.5 in each model), cCREs 2–100 kb away less so (0.58–0.64), and distal cCREs carrying the motif of the TF that the gene encodes no more than with a permuted TSS map (0.51; Fig.5f), which gave AUCs within 0.02 of 0.5. The TF–motif agreement of the shared axes therefore reflects genome-wide motif activity rather than links between TF genes and individual distal elements.

We also did an ablation study for MUTUAL’s component. We retrained the model with only its inputs changed, keeping the corpus, schedule and seeds fixed, and scored each variant on all cells of the eight datasets from tissues absent from training (Methods). Adding the foundation-model embeddings to the linear summaries never raised the error on any dataset and lowered it on three of the eight (breast, Fetal heart (ENCODE A) and Liver (ENCODE A); mean error 0.128 / 0.109 against 0.135 / 0.126 without them; Fig.5g), short of our pre-specified criterion for an improvement (lower error on at least six). The final model was likewise never worse than variants in which the foundation-model embeddings were replaced by random projections of the same width (lower error on four datasets) or taken from other cells of the same dataset (lower on two).

## 3 Discussion

MUTUAL shows that a shared RNA–ATAC space can be learned once from public multiome data and then used on new single-modality data without refitting. Its encoders are frozen single-cell foundation models joined to linear summaries of each modality, and only small networks are trained on 3.0 million paired cells. On ten datasets not used for training, this fixed model matched RNA and ATAC profiles, transferred chromatin motif activity to RNA-only cells and annotated ATAC-only cells more accurately, on average, than methods fitted on other donors of the same dataset. Because nothing is fitted to the target, the same model serves every study, and in tissues related to its training data an RNA-only or ATAC-only sample needs no multiome bridge from the same tissue [20].

The advantage was largest where the training data were most relevant. For new donors of tissues in the corpus, and for bone marrow, whose cell types are related to the blood and progenitor cells in training, MUTUAL’s matching error was 23–57% of that of the methods fitted on other donors. In tissues absent from training the gains were smaller, and MultiVI fitted on other donors matched adult heart cells better. MUTUAL is most useful when references come from other donors or studies, or when new samples must be placed without refitting. Breast, liver and heart are absent from the paired corpus, but transfer worked there, so the breadth of the paired corpus may limit generalization, but which features of a tissue determine success remains open.

What the foundation models contribute should be stated precisely. Linear summaries of expression and accessibility carried most of the information: a model without foundation-model inputs was close to the final model, and adding the foundation-model embeddings lowered the matching error on three of eight datasets. Replacing them with random projections of the same width, or with the embeddings of other cells, was worse on four and two of eight datasets and never better, so the gain comes from their content rather than from added capacity. Removing EpiAgent alone changed no dataset, whereas removing scFoundation alone raised the error on two. These results fit reports that single-cell foundation models often perform close to simple baselines when used without fine-tuning [35, 36], and suggest using them as one of several frozen views of a cell. Keeping the encoders frozen, as in vision–language models built on locked pretrained encoders [25, 37], also has a practical benefit: each network retrains in under an hour from cached features, which will make it inexpensive to update the model as new paired data appear.

The shared space carries more than cell identity. Its leading axis separated cell types in every dataset, but partners were also matched within cell types and donors far above chance, and, without any genomic coordinates, the raw view coupled genes to accessible elements near their own promoters, although removing the raw view left matching accuracy unchanged. In most datasets the shared axes also aligned the expression of TFs with the activity of their own motifs, a test defined after the first results. Most of the agreement between transferred and measured motif activity nevertheless reflects differences between cell types: a control given each query’s annotated cell type reached 0.30–0.37, against 0.37–0.44 for MUTUAL on the dataset-level readout that we adopted after the pre-specified one, and within cell types and donors the pre-specified readout reached 0.15–0.19. Predictions for individual cells within a type should therefore be read as estimates of program-level activity rather than as measurements.

The main limits of this approach concern what public data and cell-level representations can provide. First, what MUTUAL can transfer is bounded by what has been profiled in both modalities. Public multiome data are concentrated in a few tissues, in human samples and in one droplet-based assay, and cell states that no paired study has captured, such as many disease states, are placed in the shared space but receive the chromatin of their nearest profiled relatives. This boundary is not fixed: because the foundation models enter as frozen views and only small networks are trained, new paired atlases and assays, and improved foundation models, can be incorporated by recomputing features and retraining the networks. Second, the shared space describes whole cells. It carries TF-program activity and gene– promoter structure, but it does not by itself resolve which distal element regulates which gene or where a TF binds; combining cell-level transfer with sequence-based models of accessibility and with perturbation data is a natural route from programs to loci. Third, paired measurements provide ground truth only for whether a cell’s own profile is recovered. Whether transferred chromatin changes the conclusions of a new single-modality study is harder to score, and shared benchmarks of such downstream uses would help the field compare methods that connect the two layers. Within these bounds, MUTUAL turns the growing collection of public multiome data into a reusable resource that connects single-modality studies across the two layers of gene regulation.

## 4 Methods

### 4.1 Training corpus

We collected human single-nucleus multiome data (10x Genomics Multiome; RNA and ATAC from the same nucleus). Most libraries came from a search of GEO [38] through the NCBI E-utilities (31 August 2026; human series matching multiome, SHARE-seq, SNARE-seq, Paired-seq, sci-CAR, scNMT, Paired-Tag, snMultiome or joint-profiling terms; 1,078 series); heart and the studies and donors used for evaluation were excluded at intake. We added our processing of the Human Pancreas Analysis Program (HPAP) multiome data [39] and four 10x Genomics demonstration datasets. The corpus unit is a sequencing library (one donor and tissue for HPAP), called a dataset. The data span primary tissues, blood, cells derived from induced pluripotent stem cells, organoids, cell lines and xenografts, from healthy donors and patients.

The final corpus has 3.01 million cells in 400 dataset entries (183–48,956 cells per entry; median 6,753), shown in 14 tissue groups and a group of other tissues (Fig.1a). For the collection only a random 80% training split of each dataset was used. Up to 128 cells per dataset (50,007 in total) were held out to monitor convergence.

### 4.2 Preprocessing

*RNA.* From the deposited filtered count matrices (some of which had been further filtered by the authors), we kept cells with at least 100 detected genes (1,000 for HPAP) and genes detected in at least three cells, normalized counts to 10^4^ per cell, applied log(1 + *x*) and mapped genes to the 19,264-gene vocabulary of scFoundation, setting absent genes to zero.

*ATAC.* Fragments were intersected (bedtools) with the 1,355,445 hg38 candidate *cis*-regulatory elements (cCREs) that form EpiAgent’s vocabulary, [23] giving counts per cCRE; for HPAP, our peak counts were assigned to overlapping cCREs. Cells with at least 200 fragments in cCREs (1,000 for HPAP) were kept, and a cell’s EpiAgent input is its list of accessible cCREs ordered by decreasing TF–IDF weight.

*Pairing and quality.* Modalities were paired by exact barcode match among cells passing both filters; libraries with fewer than 100 paired cells were dropped. No doublet, mitochondrial-fraction or TSS-enrichment filter was applied, and 5.2% of training cells have fewer than 500 detected genes.

### 4.3 Cell-type labels

Of the training cells, 93.3% carry a cell-type label (298 label strings), used only by the NCA term of the objective. Labels of the earlier collection came from CellTypist predictions, marker-gene scoring, the original authors (one GEO study), our HPAP annotation and the 10x annotation of one demonstration dataset. The 255 libraries added later (77% of cells) were labelled automatically: each study’s cells were clustered, every cluster was named from its marker genes, starting from the 206 names of the earlier collection and adding names where needed (980 clusters, 201 names, 126 of them new), and names were propagated to cells by nearest neighbours in scFoundation feature space (Supplementary Note 1). Labels were matched by exact name; because every training batch comes from a single dataset, inconsistent naming across studies does not enter the label term.

### 4.4 Model inputs

#### Foundation-model embeddings

Both foundation models were used without updating their weights. For RNA we used the released scFoundation checkpoint [22] (sha256 9f40bf32&). For ATAC we used the released EpiAgent weights [23] (sha256 7fffb922&) and took the final-layer [CLS] output (512 values). EpiAgent reads at most 8,190 elements; for cells with more accessible elements (38% of training cells), the element list was split into interleaved subsets that were encoded separately and their [CLS] outputs averaged.

#### Linear summaries

Fitted once on a dataset-balanced sample of 199,702 training cells and then applied unchanged: for RNA, the top 256 principal components of the standardized log-expression of 4,000 panel genes; for ATAC, 256 latent semantic indexing (LSI) components [26] of the TF–IDF-weighted accessibility of the 100,000 most frequently accessible cCREs, after dropping the first component, which tracked sequencing depth.

#### Summary-view input

The foundation-model embedding and the linear summary of a cell were each scaled to unit length and concatenated 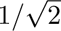 values for RNA and 768 for ATAC.

#### Raw view

The 4,000 genes with the highest variance and the 10,000 cCREs accessible in the most cells (selected on 80,000 training cells); the input is the log-normalized expression of the panel genes or the set of accessible panel cCREs.

### 4.5 Architecture

For each modality, a summary-view network (linear layer to 512 units, GELU, layer normalization, dropout 0.1, linear layer to 128 units, *ℓ*_2_ normalization) gives *z*_sum_. A raw-view network embeds each panel feature with a learned 256-dimensional vector, pools the detected features by a weighted mean (weights from relative expression for RNA and equal for ATAC, multiplied by learned per-feature weights) and applies layer normalization and a network of the same form, giving *z*_raw_. A cell’s embedding is 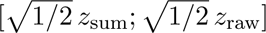, a 256-dimensional unit vector, so that the cosine similarity of an RNA and an ATAC embedding is the mean of the two views’ cosines. The model has 9.8 million trainable parameters, of which 6.2 million are used at inference (the rest are training-only decoders).

### 4.6 Training objective

Batches contain *B* = 256 cells with both modalities. With 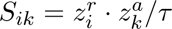 for the RNA embedding of cell *i* and the ATAC embedding of cell *k*, the loss is

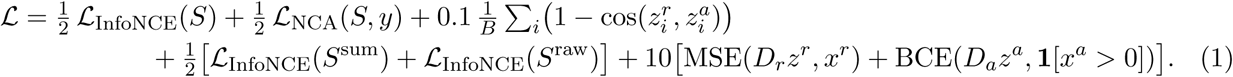

*L*_InfoNCE_ is the symmetric cross-entropy of identifying each cell’s partner among the batch in both directions. [24**?**] *L*_NCA_ is a neighbourhood-components term [27] in the multi-positive form of supervised contrastive learning: [40] for an anchor *i* with label *y_i_* and at least one other in-batch cell of that label, 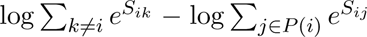, where *P* (*i*) are the other cells with label *y_i_*; it is averaged over anchors and directions, and unlabelled cells are neither anchors nor positives. *S*^sum^ and *S*^raw^ are the corresponding similarities of the two views. *D_r_* and *D_a_* are linear decoders, used only in training, that reconstruct panel-gene expression and panel-cCRE accessibility from the fused embedding. The three temperatures are learned (initialized at 0.07). During training, every detected raw-view feature of a cell is kept with probability *k ∼ U* (0.3, 1), drawn per cell; reconstruction targets are unthinned. The loss weights were inherited from earlier development rounds (Development data, below).

### 4.7 Training

Each step drew one training dataset uniformly at random and 256 of its cells, so all negatives in a batch come from the same dataset (datasets with fewer than 256 cells contributed repeated cells). Cells were drawn from a per-dataset pool of up to 2,048 cells that was refreshed as training proceeded; the schedule was precomputed and identical across model variants for a given seed. We used AdamW [41] (learning rate 10^−3^, *β* = (0.9, 0.98), weight decay 0.01) with a 500-step warm-up and a constant rate thereafter, gradient-norm clipping at 4.0 and float32 weights in PyTorch. Every model was trained for 500,000 steps. Every reported embedding comes from the snapshot at 400,000 steps, the step at which the slowest of the three main models met the convergence criterion (less than 0.5% improvement of matching error on held-out validation cells of the training datasets over the preceding 50,000 steps). Training took 49 min per model on one NVIDIA H200 GPU with all features cached in GPU memory.

We trained three models with different seeds (initialization, dropout, thinning and batch schedule). Matching (Fig.2) uses the concatenation of the three models’ embeddings, each scaled by 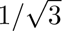, with the mean over single models beside; integration scores and annotation accuracies (Figs. 2 and 4) are means over the three models; motif transfer (Fig.3) averages the three models’ results; Fig.5 reports each model separately.

### 4.8 Embedding new data

Each modality is embedded on its own: an RNA-only cell needs only its expression profile and an ATAC-only cell only its accessible cCREs. New cells are passed through the frozen foundation models, projected onto the fixed linear summaries and panels, and embedded with the network in evaluation mode. Matching (Fig.2) uses these embeddings as they are. For transfer within a tissue (Fig.3) and for annotation (Fig.4), embeddings were centred per donor (each donor’s mean embedding subtracted per modality before renormalization, which uses the donor identities of the target data); this was applied to every method.

### 4.9 Development data and evaluation datasets

#### Development

The model’s inputs, objective and training recipe were developed over several rounds on earlier versions of the corpus, using the bone marrow, breast and liver datasets below, and the pancreatic islet and lymph node datasets; the choice of the final inputs and of the encoding of deep cells also used the ENCODE liver and heart datasets. For every result reported here, the datasets, metrics, comparators and decision rules were written down and frozen (with checksums) after the final models had been trained and before the outcomes were computed.

#### Evaluation datasets

The main evaluation panel comprised ten human multiome datasets (342,608 scored cells from 76 individuals) that were not used for training, validation or fitting of the linear summaries: breast (GEO GSE244594) [42], liver (GSE296875) [43], five ENCODE 10x Multiome datasets [44] of liver (two), fetal heart (two) and adult heart (left ventricle), bone marrow (GSE194122) [45], and pancreatic islets and pancreatic lymph nodes of HPAP donors absent from the training data [39]. The training data contain no heart, liver or breast cells. They do contain blood and haematopoietic progenitor cells, one small bone-marrow plasma-cell library (431 cells) and other HPAP donors of the islet and lymph node tissues. The fetal heart embryos came from the same tissue supplier as an atlas in EpiAgent’s pretraining data, so some donors may overlap. We scored the cells embedded by every method (for bone marrow, islet and lymph node, those passing SCARF’s cell filter), and for breast and liver fixed scoring splits whose cells are disjoint from those used to train the stored MultiVI models. Cell types came from the original authors (breast, liver), the dataset providers (bone marrow), our HPAP annotation (islet and lymph node; or RNA label transfer from the liver dataset’s labels (one ENCODE liver dataset; cross-validated accuracy 0.98); the other four ENCODE datasets have no cell-type labels, and analyses that need cell groups use clusters of RNA profiles for them (Figs. 3b and 5).

### 4.10 Comparators

Comparators were chosen so that no method sees the scored cells’ pairing, and are grouped by what they use from the evaluation dataset. *Nothing from the dataset* : SCARF [21] with its released weights (commit 9efd249) and no training. *Other donors of the same dataset* : GLUE [14] (scglue 0.4.0), MultiVI [15] and the Seurat v5 bridge integration [20] (Seurat 5.3.0). The individuals of each dataset were split into two folds balanced for cell numbers and cell types; GLUE and MultiVI were fitted on one fold (for GLUE, including its variable features, dimensionality reductions and guidance graph; for MultiVI, the paired cells of that fold’s donors) and used to embed the cells of the other fold, and every method, including ours, was scored within the held-out fold. The bridge maps single-modality cells through a bridge of paired cells, here up to 10,000 paired cells of the other fold. For matching, its reference was the RNA of all cells of the other fold; the scored fold’s ATAC profiles were mapped through the bridge, and its RNA profiles were mapped to the reference by Seurat reference mapping and then transformed as the bridge transforms reference cells, so that both modalities lie in the reference’s bridge representation (this RNA projection is ours; it reproduces exactly the representation Seurat computes for the reference cells). The bridge was run on the datasets with cell-type labels (six of the main panel). CelLink [18] matched each query to its reference by optimal transport with settings fixed before the results were computed. For annotation, the bridge’s reference was the labeled RNA of the other donors of the query’s fold, with the same bridge cells, and we also ran scJoint [16] (official code, commit cbbfa5d; trained on the labeled RNA of the other donors of the query’s fold and the ATAC profiles of the other fold, with the query embedded afterwards) and a gene-activity nearest-neighbour baseline that uses no trained model.

### 4.11 Cross-modal matching

In a pool of *m* cells measured in both modalities, the FOSCTTM [28] of an RNA query is the fraction of the *m−* 1 other cells whose ATAC embedding is more similar (cosine) to the query than the query’s own ATAC embedding; ATAC queries are scored symmetrically. Lower is better and 0.5 is chance. The two directions are reported separately. Pools were the whole dataset (dataset scope, primary), each donor’s cells (donor scope, co-primary) or one donor’s cells of one type (within-type scope; at least 64 cells, subsampled to at most 2,000). A dataset’s value is the cell-weighted mean over its pools. For the comparison with the methods that use other donors, every method was scored within each of the two folds: at the dataset scope the pool was the cells of one fold and the two folds’ values were averaged, and donor and within-type pools, which lie within one fold, were averaged over both folds with cell weights. Panel means weight datasets equally. In each direction, MUTUAL was called better (worse) than a comparator on a dataset if its error was lower (higher) by at least 0.005 and by at least twice the standard deviation of the errors of its three single models; a dataset call requires the same call in both directions and is otherwise reported as mixed, or as no difference when neither direction differs. The rule was pre-specified for GLUE and MultiVI at the dataset and donor scopes and, when the bridge was added, for the bridge at the dataset scope; comparisons with SCARF and at the within-type scope are descriptive.

### 4.12 Integration scores

For each method, a cell’s RNA and ATAC embeddings were scaled to unit length and concatenated (the single-modality inputs, scFoundation, EpiAgent, PCA and LSI, were scored on their own), and the scIB metrics [29] (scib 1.1.7) were computed on at most 4,000 labeled scored cells per dataset (the same cells for every method): normalized mutual information and adjusted Rand index at the best of 19 Leiden resolutions [46] (0.01–2.0; five seeds), cell-type silhouette and cLISI, and batch silhouette, iLISI and graph connectivity with donor as batch; overall = 0.6*×* conservation + 0.4*×* mixing. Isolated-label scores were not used. Because each cell’s two embeddings are concatenated, these scores assess the joint representation of paired cells. MUTUAL’s scores are means over its three models, and it was called better than another representation on a dataset if its overall score was higher by at least 0.01 and by at least twice the standard deviation across its three models. GLUE and MultiVI were scored as fitted on all cells of each dataset. For Fig.2d, UMAP [47] (umap-learn; 30 neighbours, minimum distance 0.3, cosine distance) was computed on the joint embedding of all 18,305 bone-marrow cells, with each modality represented by the concatenated embeddings of the three models.

### 4.13 Transfer of chromatin TF-motif activity to RNA-only cells

#### Measured activity

For each evaluation dataset, chromVAR [10] deviation scores were computed from each cell’s own ATAC profile with a Python re-implementation (50 background peak sets matched for GC content and accessibility) on the cCREs accessible in at least 25 cells, for the 765 JASPAR 2022 [48] CORE vertebrate motifs with at least 50 hits (MOODS [49] scan, *P <* 10^−4^, 500-bp windows centered on each cCRE).

#### Transfer and accuracy

An RNA-only query received the softmax-weighted mean (similarity/0.05) of the measured deviations of its 20 nearest reference ATAC cells; every co-embedding method was evaluated through this procedure with the same queries and references. Agreement was scored over all scored query cells of a dataset (in the fold design, of a fold), after regressing quality covariates out of transferred and measured scores across those cells; the reported accuracy is the mean Pearson correlation over the 100 motifs with the largest variance of measured activity across those cells (the 100 motifs of the within-stratum readout give similar values). A cell-type-only control gave each query the mean measured deviations of the reference cells of its own cell type (Leiden clusters of RNA profiles for the four datasets without labels). This dataset-level readout was adopted after the results of the pre-specified readout had been seen; the pre-specified readout scored agreement within strata of donor *×* cell type with at least 30 query cells, after regressing quality covariates out within each stratum, over the 100 motifs with the most within-stratum variation, and is reported in Extended Data. Shuffled controls gave *|r| <* 0.001. Because the dataset-level readout spans donors, a prediction that carries only the reference donors’ mean activity correlates negatively with the query donor’s own mean; a reference-permuted control was therefore negative (*−*0.03 on average), and this term, which no reference-based transfer can remove, lowers every method’s agreement in the cross-donor settings.

#### Settings and comparators

References were the ATAC profiles of other donors of the same dataset (all, or random subsets of 100, 300 or 1,000 cells; five draws per size and query donor), the ATAC halves of a random half-split within donor *×* cell type (same-donor setting), or 2,500 cells from each dataset of another tissue (cross-tissue). For the comparison with GLUE and MultiVI fitted on other donors (Fig.3b), queries and references were drawn from the same fold of donors, GLUE and MultiVI embedded that fold with the model fitted on the other fold, and breast, which has too few cells, has no 1,000-cell level. Embeddings were centered per donor before the neighbor search. Comparators were GLUE, MultiVI and SCARF through the same procedure and CelLink; TF activity estimated from RNA alone with decoupleR [9] (1.9.2; CollecTRI network [50]) or from the TF’s own mRNA; and, as a reference that uses paired cells of the target dataset, a ridge regression [51] from the reference cells’ paired RNA to their measured activity. Intervals are 95% donor-block bootstrap intervals (1,000 resamples). On a dataset, a method was better than another at subsampled references if its mean advantage was at least 0.010 and positive in all five draws, and with full references if the advantage was at least 0.010 with an interval excluding zero; a pre-specified secondary view excluded AP-1 and other bZIP motifs.

### 4.14 Stem-cell-derived islet use case

We used the multiome data of stem-cell-derived islets of ref. 30 (GEO GSE199636), all of which were excluded from the training, validation and fitting of the model used here: the 14 libraries of the maturation series (97,247 cells), namely baseline SC-islets (2 libraries), SC-islets after 2, 3 and 4 weeks and 6 and 12 months in stage 6 of the differentiation protocol in culture (5), grafts recovered after transplantation into mice (3) and primary human islets (4), as described in the GEO records. Cell types were transferred by RNA nearest neighbors from the authors’ labeled cells of the baseline and primary libraries (cross-validated accuracy 0.90).

#### Trajectory

We used the *β* cells (23,576 of 29,519 cells pass our corpus quality control; per-library medians of their maturation score differ from those of the excluded cells by at most 0.03) to build and analyze the trajectory: UMAP (30 neighbours, minimum distance 0.3, cosine, no PCA) of the MUTUAL RNA embedding (three-model ensemble), then Monocle3 [31] (1.4.26, R 4.4.3; cluster cells on the UMAP, then learn graph). The readouts, fixed before the run, were the Spearman correlation between library medians of pseudotime and stage order. The default fit (partitions, one root per partition at the node richest in baseline cells) placed primary islets in a separate partition. After seeing it, we report a single graph over all cells (use partition = FALSE, closed loops) rooted at the tip of the graph (a node of degree one with at least ten assigned cells) whose cells are the most often baseline cells. Because re-fitting did not reproduce the graph exactly (cluster cells varied between runs), the pseudotime for this root was computed on the fitted graph as Monocle3 defines it, as the geodesic distance along the graph from the root to each cell’s projection onto its nearest edge; with Monocle3’s own root this reproduces its pseudotime (Pearson *r* = 0.9996).

#### Motif activity by stage

Every RNA profile of the series received the softmax-weighted mean (similarity/0.05) chromVAR deviations of its 20 nearest cells, by exact cosine search over all cells of one of two ATAC banks: the ATAC profiles of the other libraries of the series (each query’s own library excluded), and the 3,009,609 ATAC profiles of the training corpus, which contains no stem-cell-derived cells. For the corpus bank, the chromVAR expectation and background peak sets were defined once from all corpus cells on the same cCRE universe and motif set as the measured activity of the series, so that each bank cell’s deviations do not depend on which cells are retrieved. Before any prediction was computed, we fixed the cell set (*β* cells), the aggregation (library means, then stage means), the consistency statistics (per-motif Pearson correlation of predicted and measured library means across the 14 libraries, and agreement in the direction of the primary-versus-baseline, month-12-versus-baseline and graft-versus-culture differences) and a list of 40 TF motifs: 36 whose change between these stages is reported in the text, figures or supplementary tables of ref. 30 (with the reported direction). A data-driven list of the 30 motifs that best separate the 14 libraries in the measured activity (largest *F* statistic, pruned at *|r| >* 0.9) was analyzed in the same way. With one library per time point in culture, these comparisons are descriptive.

### 4.15 Annotation of ATAC-only cells

In each labeled dataset, every donor was held out in turn: its ATAC profiles were the queries and the labeled RNA of other donors the reference. In the primary, fold design (Fig.4b,c), the reference was the other donors of the query’s fold, every method that is fitted to target data was fitted on the other fold only, and every method used the same references; in the all-donor reference design (Fig.4d, the novel-type analysis and the fits on all cells), the reference was all other donors. For bone marrow the held-out unit was a sample (one donor at one site, or one eye), so other samples of the same individual can be in the reference. Embeddings were centered per donor and scaled to unit length; each query took a similarity-weighted vote (softmax of cosine similarity, temperature 0.05) of its 15 most similar reference cells, the winning share serving as confidence. Queries whose type had at least five reference cells were scored; accuracy was the balanced accuracy pooled over held-out donors, with 95% intervals from 1,000 donor-block bootstrap resamples, and a method was called more or less accurate than another on a dataset if the interval of the paired difference excluded zero. Accuracy–coverage curves ranked queries by confidence. To test novel types, each type with at least 100 cells was removed from the reference in turn and its cells were scored by their mean similarity to the 15 nearest remaining reference cells. Permuted reference labels gave chance accuracy.

### 4.16 What the shared space aligns

Each of the three models was analyzed separately on uncentered embeddings. Within each dataset, cells were split in half within donor *×* cell group (cell groups: author labels, or clusters of RNA profiles for the four ENCODE datasets without labels). On one half we computed the eigenvectors of the symmetrized cross-covariance between RNA and ATAC embeddings (shared axes); on the other half we measured the share of the mean partner similarity carried by each axis, the number of leading axes that recover 90% of the matching gain over chance (the reduction in matching error from 0.5 to that of the full embedding) and the share of the leading axis’s variation that lies between cell groups (an identity axis if at least 0.5 in all three models). Exact Shapley values [32] attributed the cross-modal similarity to classes of cell covariates: cell group, donor *×* cell group, read depth, quality metrics, cell-state programs and TF programs (the first 16 principal components of TF-motif activity and of TF mRNA, within donors). Matching within cell groups used pools of 32 cells of one cell group and donor, against a permutation floor and a depth-only scorer that pairs cells by their numbers of detected panel genes and accessible panel cCREs.

For TF programs, we computed within donors the correlation of each axis’s ATAC-side score with each motif’s chromVAR deviation and of its RNA-side score with each TF’s mRNA (TFs detected in at least 5% of cells; motifs matched to TFs by name), averaged them within JASPAR families and correlated the family means (Spearman). The pre-specified test compared this correlation on each of the five leading axes with 10,000 permutations (Benjamini–Hochberg adjustment over the five axes) and was informative only where its positive control, one-versus-rest contrasts between cell groups scored with the same statistic, was detected. Because its within-donor negative control gave false positives in five datasets, we adopted after the first results had been seen a comparison with random shared axes from 40 within-donor permutations per axis: the correlation, with one TF per family, pooled over the five leading axes was compared with 10,000 resampled combinations of these permutations, across cells and within donor *×* cell group; a dataset counts if *P <* 0.01 in at least two of three models. For the islet example (model 1), cell types with at least 20 cells were placed at their mean score on the leading axis (average of the RNA-side and ATAC-side scores), genes were ranked by the within-donor Pearson correlation of their expression with the RNA-side score (genes detected in at least 1% of cells), and the family means of all 65 TF families are shown. Finally, we measured from gradients of the cross-modal similarity how strongly the raw view couples each panel gene to each panel cCRE, and compared cCREs near the gene’s own transcription start site (GENCODE v48 [52]) with matched control elements.

### 4.17 Input ablations

Variants differed from the final model only in their inputs, with the same corpus, panels, schedules, seeds and read step: scFoundation or EpiAgent alone with the other modality’s linear summary, both foundation-model embeddings without summaries, and the summaries without foundation-model inputs (NF-LSI); variants without the raw view (which kept the reconstruction term); a content control, in which each cell’s scFoundation and EpiAgent embeddings were taken from two other cells of the same dataset (also at test time); a width control, in which the foundation-model embeddings were replaced by fixed Gaussian random projections of the panel-gene expression (to 3,072 values) and of the TF–IDF accessibility (to 512 values); and the final model with the EpiAgent or the scFoundation embedding removed from its summary view. Variants were compared on dataset-scope (primary) and donor-scope (co-primary) matching error over the datasets from tissues absent from training, with islet and lymph node as guards. On a dataset, one variant was better than another if its seed-mean error was lower by at least 0.005 and by at least twice the larger seed standard deviation, in both directions. Pre-specified summary labels were: helps (better on at least six of nine datasets), complementary (better on at least six and worse on none) and non-inferior (worse on none); we report the counts over the eight datasets of the main panel, which give the same labels.

## Notes

### Competing Interest Statement

S.C. is the co-founder and shareholder of iOrganBio. The other authors have no conflict of interest.

### Funding

- National Institutes of Health: OT2OD038003, U24DK138515

